# Seasonal physiological responses to heat in an alpine range-restricted bird: the Cape Rockjumper

**DOI:** 10.1101/248070

**Authors:** Krista N Oswald, Alan TK Lee, Ben Smit

## Abstract

Hot, dry summer conditions impose physiological stress on endotherms, yet we have a poor understanding of how endotherms seasonally adjust their costs of thermoregulation under hot conditions. We determined whether seasonal phenotypic plasticity in evaporative cooling capacity at high temperatures explained how the range-restricted Cape Rockjumper (*Chaetops frenatus*; hereafter “Rockjumper”), copes with hot and dry summer temperatures of the temperate mountain peaks of southwest South Africa. We measured evaporative water loss (EWL), resting metabolic rate (RMR), and body temperature (T_b_) at high air temperatures (30 to 42°C) of individuals from a wild population of Rockjumpers during winter and summer (n = 11 winter, 4 females, 7 males; n = 10 summer, 6 females, 4 males). We found Rockjumper evaporative cooling in summer imposes higher EWL (i.e. greater water costs) compared to winter, although an accompanying lack of change in RMR resulted in increased summer cooling efficiency. These patterns are similar to those observed in species that inhabit regions where summer temperatures are routinely hot but not water stressed. Our findings show that avian seasonal physiological adjustments to heat can be diverse. Further seasonal studies on thermoregulation in the heat will greatly improve our knowledge of the functional value traits such as evaporative cooling efficiency and heat tolerance hold and how they contribute to the physiological stress organisms experience in heterogenous environments.

## Introduction

Many birds live in temporally heterogenous environments where physiological phenotypes are adjusted seasonally to optimise the balance of energy (typically heat and chemical energy) and water. Birds provide useful model systems to study energy and water balance as they are generally small, have high body temperature, and very high mass-specific rates of metabolic heat production. Whereas many studies to date have focussed on physiological adjustments to winter conditions (Swanson 1991, Swanson 2010, McKechnie et al. 2015), studies about adjustments to hot summer conditions are rare. Hotter summer conditions will impose physiological stress on endotherms through greater rates of environmental heat gain, decreased rates of metabolic heat dissipation, and often elevated evaporative water loss (EWL) rates (Wolf 2000). Loss of body water due to higher levels of EWL cannot be sustained indefinitely without risking dehydration (Tieleman et al. 2002), so presumably there are strong selective pressures to minimize the costs of heat dissipation, especially in water scarce environments.

Birds are often exposed to high operative temperatures (defined as an integrated temperature meant to reflect the overall thermal environment experienced by individuals: as most are diurnal and forage in exposed terrestrial environments (e.g., Wolf 2000, Tieleman 2002, Williams and Tieleman 2005). Evaporative water loss is well documented as the primary physiological mechanism of heat dissipation in birds when air temperature (T_air_) exceeds normal body temperature (T_b_). Elevated water demands may therefore represent a primary thermoregulatory cost during summer.

Our current understanding of the thermoregulatory costs associated with summer acclimatization to higher temperature relies on a handful of acclimation (typically exposure to hot laboratory holding conditions) and acclimatisation (adjusted to natural weather patterns in free-ranging and captive birds in outdoor holding conditions) studies. These acclimation studies have often demonstrated that heat responses involve an increase in evaporative cooling efficiency (defined as the ratio of evaporative heat lost to metabolic heat produced) to better control T_b_ when exposed to hot conditions (Marder and Arieli 1988, Ophir et al. 2002, Tieleman and Williams 2002, McKechnie and Wolf 2004). This generally involves an ability to elevate total EWL rates (McKechnie and Wolf 2004), and/or reduce RMR (resting metabolic rate) at hot temperatures (Tieleman and Williams 2002) in order to regulate body heat. Thus, T_b_ can be maintained by elevating EWL at high T_air_ to dissipate metabolically produced heat, and/or reducing RMR to decrease the initial amount of metabolically produced heat. Alternatively, despite the potentially lethal risks of hyperthermia, birds may use some degree of facultative hyperthermia, allowing T_b_ to rise, presumably to promote savings in EWL (Tieleman and Williams 1999). We may expect facultative hyperthermia to be more beneficial in water scarce environments, but it seems to be widely used by birds and the costs and benefits are not yet clear (Tieleman and Williams 1999, Williams and Tieleman 2005).

Seasonal studies on thermoregulation in the heat have thus far shown summer-acclimatized birds to generally increase evaporative cooling efficiency leading to improved regulation of normal T_b._ For example, captive-bred Houbara Bustards (*Chlamydotis macqueenii*) achieved greater evaporative cooling capacity during summer by increasing total EWL and decreasing RMR (Tieleman et al. 2002). In contrast, free-ranging Freckled Nightjars (*Caprimulgus tristigma*) also achieved greater cooling capacity, but solely through elevated total EWL rates (O’Connor 1995). The mechanisms of seasonal variation in evaporative efficiency can also vary at an intra-specific level, with Noakes et al. (2016) showing that free-ranging White-browed Sparrow-weavers (*Plocepassar mahali*) from mesic sites increased total EWL and decreased RMR. The emerging patterns from these studies suggest that seasonal variation in physiological stress and costs of evaporative cooling may vary at both intra- and inter-specific levels with local climate.

To date, the handful of studies reporting seasonal responses to high temperatures have focussed mostly on wide-ranging, predominantly desert species that routinely experience hot summer conditions across their range (e.g., Noakes et al. 2016, O’Connor et al. 2016). To better understand the role of seasonal responses in evaporative cooling capacity at reducing physiological stress to warming conditions, we also need data from species that inhabit dynamic climates where summers can be 60 to 80 °C warmer than winter (e.g. high-latitude continental climates), as well as seasonally stable climates where mean temperatures vary by less than 10°C (e.g. some Mediterranean, warm temperate or cool tropical climates).

Our purpose was to assess seasonal adjustments in T_b_, EWL, RMR, and evaporative cooling efficiency at high air temperatures (i.e. T_air_ > 30 °C) in a 50-g passerine bird, the Cape Rockjumper (*Chaetops frenatus*; hereafter “Rockjumper”), mostly restricted to high-elevation Cape Fold Mountains of South Africa. Rockjumpers are an interesting species to assess seasonal responses to heat as annual mean air temperatures across their range are generally mild [14.3 ± 1.6 °C; (Lee and Barnard 2015b)], and climate warming combined with their restricted range may require Rockjumpers to cope with higher air temperatures than previously experienced. While these mountains will periodically experience T_air_s that approach or exceed 40°C when heat waves precede the landfall of mid-latitude frontal systems (Mucina and Rutherford 2006), these events are rare, and the preferred mountain habitat of Rockjumpers has historically been stable (Cowling et al. 2015). Thus, whether Rockjumpers possess the ability to cope with hot temperatures above their proposed thermal niche may reveal how cool-climate species may adapt to climate warming.

Since Cape Fold Mountain summers are typically dry, we hypothesise that summer adjustments in EWL and T_b_ patterns centred on water conservation (lower summer EWL, and lower RMR) may have incurred a selective advantage in Rockjumpers. In addition, given the relatively mild climate of their habitat, we predict that Rockjumpers may show a limited capacity to improve evaporative efficiency during summer, compared to desert birds’ species studied thus far.

## Methods

### Study Site and Species

This study took place at Blue Hill Nature Reserve (BHNR; 33.59 S; 23.41 E; 1000 – 1530 m, above sea level) in the Western Cape Province, South Africa. Rockjumpers occur in the highest ridges and slopes of the reserve, always at very low densities. The specific territories occupied by Rockjumpers used in this study ranged from ~1100 – 1500 m above sea level. Rainfall and temperature data were collected every 30 minutes using an on-site weather station (Vantage Vue, Davis Instruments Corp., USA). Average daily temperature from two weeks before the first bird was captured until the day the last bird was captured was 8.7 °C ± 6.0 in winter (July 24 - August 31 2015) and 20.5 °C ± 5.9 in summer (January 1 - 31 2016). Temperature minima and maxima occurring over the study period were -2.6 °C and 27.5 °C in winter, and 6.9 °C and 35.4 °C in summer. Average dew point temperatures (a good proxy of absolute humidity) over a three-year period (2013 to 2015) at BHNR are 1.9 °C for July and 12.1 °C for January (A.T.K.L. unpublished data).

During July and August 2015 (winter) 16 individuals were captured (nine males, seven females), and during January 2016 (summer) 11 individuals were captured (five males, six females). Body mass (M_b_) was measured to within 0.1 g before and after each experimental procedure with all birds maintaining mass within 5 % capture mass (average M_b_ (g): winter = 54.4 ± 3.2, summer = 52.3 ± 3.1). We released one male in winter (due to M_b_ loss of > 5 %) and one female in summer (prolonged agitation at T_air_ > 39 °C) before experimentation, making the sample size 15 in winter (eight males, seven females) and 10 in summer (four males, six females). After all experimental runs birds were weighed and returned to holding cages and provided with tenebrionid beetle larvae *ad libitum*. Birds were kept in captivity for a maximum 48 h period. We measured physiological responses to heat (current study), and physiological responses to cold and basal metabolism (unpublished data), in each individual during the captive period. We allowed birds to rest in holding cages for at least five hours between experiments. Measurements at high temperature therefore occurred either on day of capture (if caught before 15:00) or day after capture (if caught after 15:00); the exact time of measurement depended on the number of birds caught on a given day (n = 1-4), with 1-3 experimental heat runs per day.

### Body Temperature Measurements

Individual birds were injected with a small, temperature-sensitive, Passive Integrated Transponder (PIT) tag intra-peritoneally to measure T_b_ throughout our experiments. PIT-tags provide T_b_ information while minimising handling effects, with no significant alteration of individual condition (Gerson et al. 2014, Ratnayake et al. 2014), and no significant negative long term effects on free-ranging individuals (Ratnayake et al. 2014) or amongst Rockjumpers specifically (Oswald et al. 2017).

### Metabolic Measurements

We followed Whitfield et al. (2015) by measuring metabolic rate, evaporative water loss and body temperature in birds over a few hours while being exposed to a ramped air temperature profile. Birds typically spent between two and 24 hours in captivity before physiological responses were measured at high temperatures. Birds were placed individually in a 4-L airtight plastic chamber (Lock & Lock, India) fitted with a wire-mesh platform raised 15 cm from the floor to ensure normal perching posture. To determine that water vapour absorption within the plastic respirometry chambers were negligible, we followed Whitfield et al. (2015) by comparing rates of change in CO_2_ and water vapour by switching air streams that varied considerably in CO_2_ and water vapour. A thin layer of mineral oil was used to ensure faecal water did not factor into EWL measurements. Chamber temperature (T_air_) was measured using a thermistor probe (model TC100, Sable Systems, USA) inserted one cm through a small hole in the lid.

Respirometry chambers were placed in a custom-made environmental chamber consisting of a100-L cooler box lined with copper tubing through which temperature-controlled water was pumped from a circulating water bath (FRB22D, Lasec, South Africa). A small fan was placed inside the 100-L environmental chamber to ensure a uniform distribution of air temperature. Continuous monitoring of bird stress was provided by an infrared light source and closed-circuit security camera with live video feed. A BioMark PIT tag reader was placed next to the chamber to record T_b_ every minute (Gerson et al. 2014, Whitfield et al. 2015).

Birds were habituated to the chamber for between 10 to 15min before the experiment started. This period of acclimation to the respirometry chamber was shorter than in the study of Gibbons and Andrews (2004), as we aimed to shorten the time Rockjumpers were exposed to hot conditions. We observed that birds were very calm upon placement in to the respirometry chambers and that gas traces typically stabilised within the first 10 minutes in the chamber. After the acclimation period, EWL and RMR (measured indirectly as oxygen consumption (V□_O2_) and carbon dioxide emission (V□_CO2_)) were measured in mL min^-1^ using a portable open-flow respirometry system. For all measurements, flow rate of atmospheric air through bird chambers was controlled using FMA-series mass flow controllers (Omega, USA) calibrated using a 1-L soap bubble metre (Baker and Pouchot 1983). Atmospheric air was supplied at flow rates around 3 L min^-1^ (occasionally as low as 1.75 L min^-1^) to ensure [O_2_] to the chamber remained within 0.5% of incurrent [O_2_]. This allowed 95% wash-out rates, calculated using the corrected equation 8.1 in Lighton (2008), of around 4 minutes at our maximum flow rates. Our air pump did not allow for flow rates greater than 3 L min^-1^, and chamber humidity levels were thus higher than those used by Whitfield et al. (2015) for desert passerines. However, our aims were not to determine thermal tolerance limits as in the latter study, but rather to test thermoregulatory responses similar to the extreme maxima T_air_ and humidity levels these birds are likely to experience in their natural environment.. Atmospheric air was scrubbed of water vapour using columns of drierite, so that water vapour free air entered the bird chamber. Chamber humidity levels were maintained at water vapour levels of 10 ppt (range 6 and 14 ppt), equivalent to an average dew point temperature of around 8 °C. The maximum levels of humidity that the Rockjumpers experienced in the chamber were low enough to ensure high water vapour pressure deficits and effective evaporative cooling (see (Gerson et al. 2014). Subsampled air was then pulled from the bird chamber through a water vapour analyser (RH-300, Sable Systems, USA) before entering O_2_ and CO_2_ analysers (Foxbox-C Field Gas Analysis System, Sable Systems, USA). The Foxbox included a subsampling pump, and allowed for analog outputs to be digitized and recorded at one-second intervals using Expedata Data Acquisition and Analysis Software (Sable Systems, USA).

### Experimental Protocol

Birds were subjected to a ramped series of T_air_ (30, 33, 36, 39, 42 °C) for 15 to 20 minutes each, following Whitfield et al. (2015). Although these air temperatures are higher than Rockjumpers likely experience in their natural environment, operative temperatures can be 10 to 20 °C above air temperature when directly exposed to the sun and/or near the soil surface (Tieleman 2002). Moreover, the maximum test T_air_ of around 42 °C allowed that we could determine thermoregulatory responses at T_air_s above expected normal avian T_b_ (~41.6 °C (Prinzinger et al. 1991) where evaporative cooling demands become a necessity for maintaining heat balance (Wolf 2000). Baseline values of O_2_, CO_2_, and water vapour pressure, were recorded for a minimum of five minutes at the beginning of each experimental test, as well as between each T_air_ and again at the end of the run. Birds were held at the final T_air_ of 42 °C for up to 20 minutes to allow a thorough assessment of RMR, EWL, and T_b_ regulation at T_air_ close to T_b_ (Marras et al. 2015, Whitfield et al. 2015). Heat tolerance measurements were taken during the active phase of birds within 48 hours of capture. Birds were in chambers for a mean ± SD of 106.5 ± 12.9 minutes, with no birds kept in the chamber more than 130 minutes.

Expedata data files were corrected for O_2_ drift in baselines using the relevant algorithms in Warthog LabHelper (http://www.warthog.ucr.edu). All measurements were taken as the minimum V□_O2_ over a 60 second interval during the last five minutes at each T_air_ for calm birds to ensure the most accurate measurements of RMR. We calculated total metabolic rates, as it has been suggested they are more informative than mass-specific values for seasonal comparisons (Swanson 1991, Cooper 2002). Outgoing flow rate and rates of V□_O2,_ V□_CO2_, and V□_H2O_ were calculated using equations 9.3, 9.4 and 9.4, and 9.6 from Lighton (2008) respectively. RMR values are presented in Watts (W; also representative of metabolic heat production, MHP) and calculated using a Joule conversion of 20.1 J mL^-1^ O_2_ (Walsberg and Wolf 1995). We calculated an average respiratory quotient of 0.76 using our measured V□_O2_ and V□_CO2_ values (winter average = 0.76 ± 0.01, summer average = 0.76 ± 0.10). Evaporative heat loss (EHL, W) was calculated from EWL using latent heat of vaporisation 2.4 J mg^-1^, and evaporative cooling efficiency was calculated as the ratio of EHL to MHP.

### Statistical Analysis

We removed four individuals from our winter sample due to signs of breeding (i.e. brood patches; one male, two females), leaving a sample size of 11 in winter (seven males, four females). To explore the contribution of potential predictor variables (i.e. M_b_, sex, T_air_, and season) on response parameters (RMR, EWL, T_b_, EHL/MHP), linear mixed-effects models were fitted using the *lme4* package (Bates et al. 2013) in R version 3.1.2 (The R Foundation for Statistical Computing, 2014), with individual included as a random effect (see Supplementary file “Table S1” for raw data). We used the ANOVA function from the *car* package (Fox et al. 2007) to examine models for significant parameters as suggested by Zuur et al. (2009).

For all parameters, significance did not vary with whole-animal vs. mass-specific values, so we only report whole-animal values. We created a base model for each response parameter that was a function of the interaction between T_air_ and season, including T_b_, sex, and mass as covariates. Best models were found by sorting through a list of competing models created from the dredge function using *MuMIn* (Barton 2013) based on AICc (corrected AIC for small sample size). We present results for all model within 2Δ AICc of the top model. For all parameters, sex and mass were not included in best models.

For each heat response parameter data spread was inspected in response to increasing T_air_ for possible inflection points in the response variable, as significant inflection points were found for similar heat parameters in Milne et al. (2015). Where inflection was suspected (EWL, EHL/MHP, T_b_), the breakpoint value was assessed using the segmented function in *segmented* (Muggeo 2008) by testing linear models of the parameter of interest as a function of T_air_ and season, with multiple parameters for inflection point possibilities, selecting the best model by AICc. Values are presented as mean ± standard error (SE) except where indicated otherwise.

## Results

### Evaporative Cooling Efficiency

The estimated EHL/MHP inflection point was 37.3 ± 0.8 °C in summer and 33.5 ± 0.8 °C in winter. Above these inflection points EHL/MHP increased significantly with increasing T_air_ (mean slope ±SD: summer = 0.05 ± 0.01 EHL/MHP °C^-1^, winter = 0.03 ± 0.00 EHL/MHP °C^-1^; χ^2^_1,50_ = 83.43; p < 0.01) and the interactions of T_air_*season was significantly different between seasons (χ^2^_1,50_ = 23.95; p < 0.01; Fig 1; see supplementary materials Table S2). There was a seasonal effect for EHL/MHP, with EHL/MHP higher in summer compared to winter for the experimental range of T_air_ (mean EHL/MHP ± SD: summer = 0.4 ± 0.14, winter = 0.28 ± 0.1), as well EHL/MHP higher above average T_b_ (i.e. T_air_ > 40 °C) in summer compared to winter (mean EHL/MHP ± SD: summer = 0.79 ± 0.09, winter = 0.69 ± 0.03).

**Figure 1.**
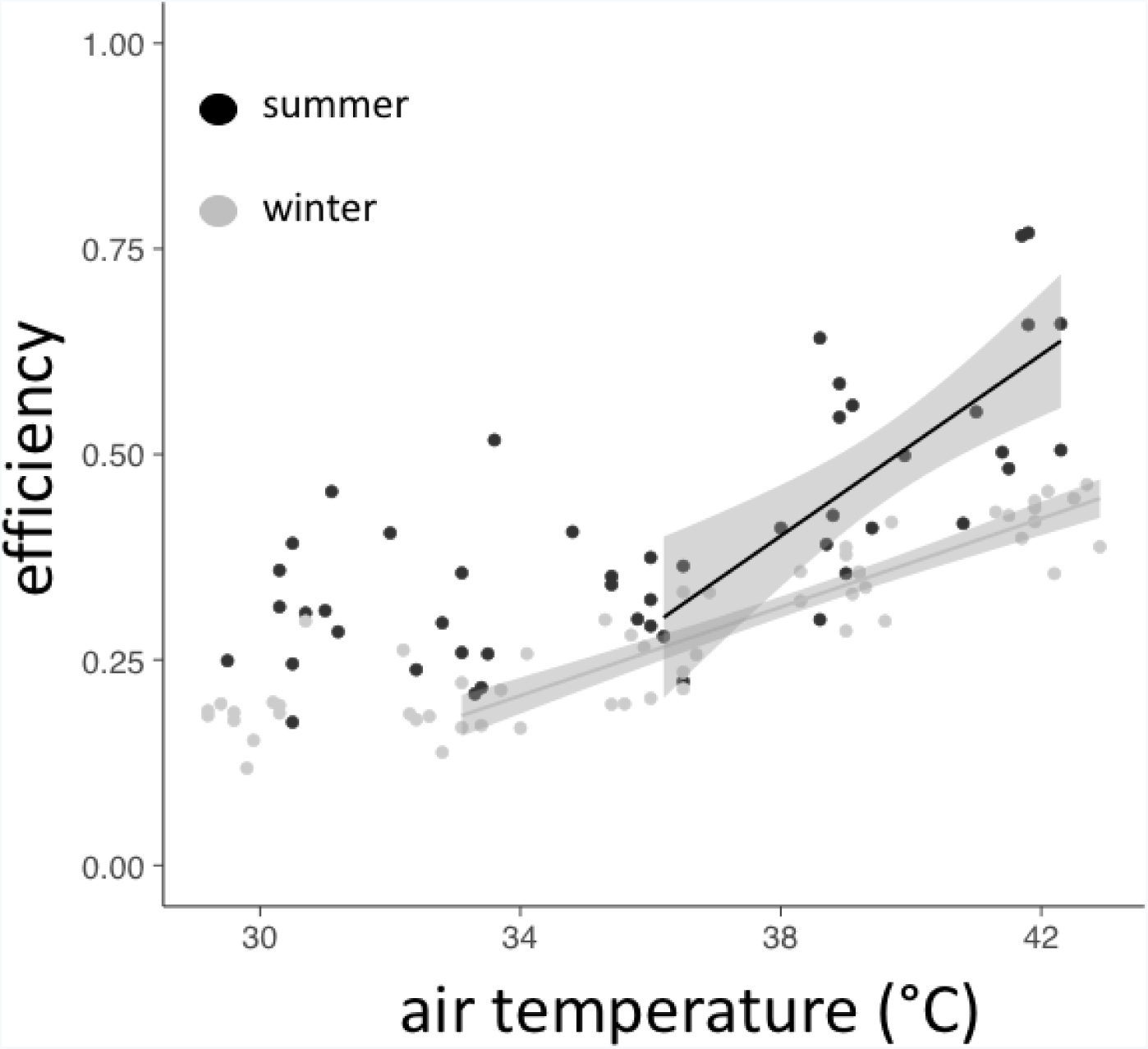
Evaporative cooling efficiency (evaporative heat loss/metabolic heat production) data over a range of air temperatures (29 – 43 °C) collected for Cape Rockjumpers (*Chaetops frenatus*) captured in summer (n = 10) and winter (n = 11) at Blue Hill Nature Reserve, South Africa. Significant difference was found above an inflection point of 36.2 °C, with separate seasonal trendlines above inflection points and 95% CI shown.

### Evaporative Water Loss

Summer improvements in evaporative cooling efficiency were corroborated by increased EWL at high T_air_ during summer. The estimated EWL inflection point was 36.0 ± 1.6 °C in summer and 34.7 ± 0.9 °C in winter. Above inflection points EWL (mg hr^-1^) increased significantly with increasing T_air_ (mean slope ± SD: summer = 90.41 ± 14.20 mg hr^-1^ °C^-1^, winter = 49.82 ± 3.66 mg hr^-1 °^C^-1^; χ^2^_2,67_ = 97.02; p < 0.01), with a notable difference in seasons: the summer mean EWL was significantly higher in this temperature range (mean EWL ± SD: summer = 783 ± 397 mg hr^-1^, winter = 447 ± 166 mg hr^-1^, χ^2^_2,67_ = 9.75; p < 0.01;; see supplementary materials Table S3). The rate of change for EWL (mg hr^_1^) was also significantly greater in summer compared to winter (χ^2^2,67 = 0.00; p < 0.01; see supplementary materials Table S3).

**Figure 2.**
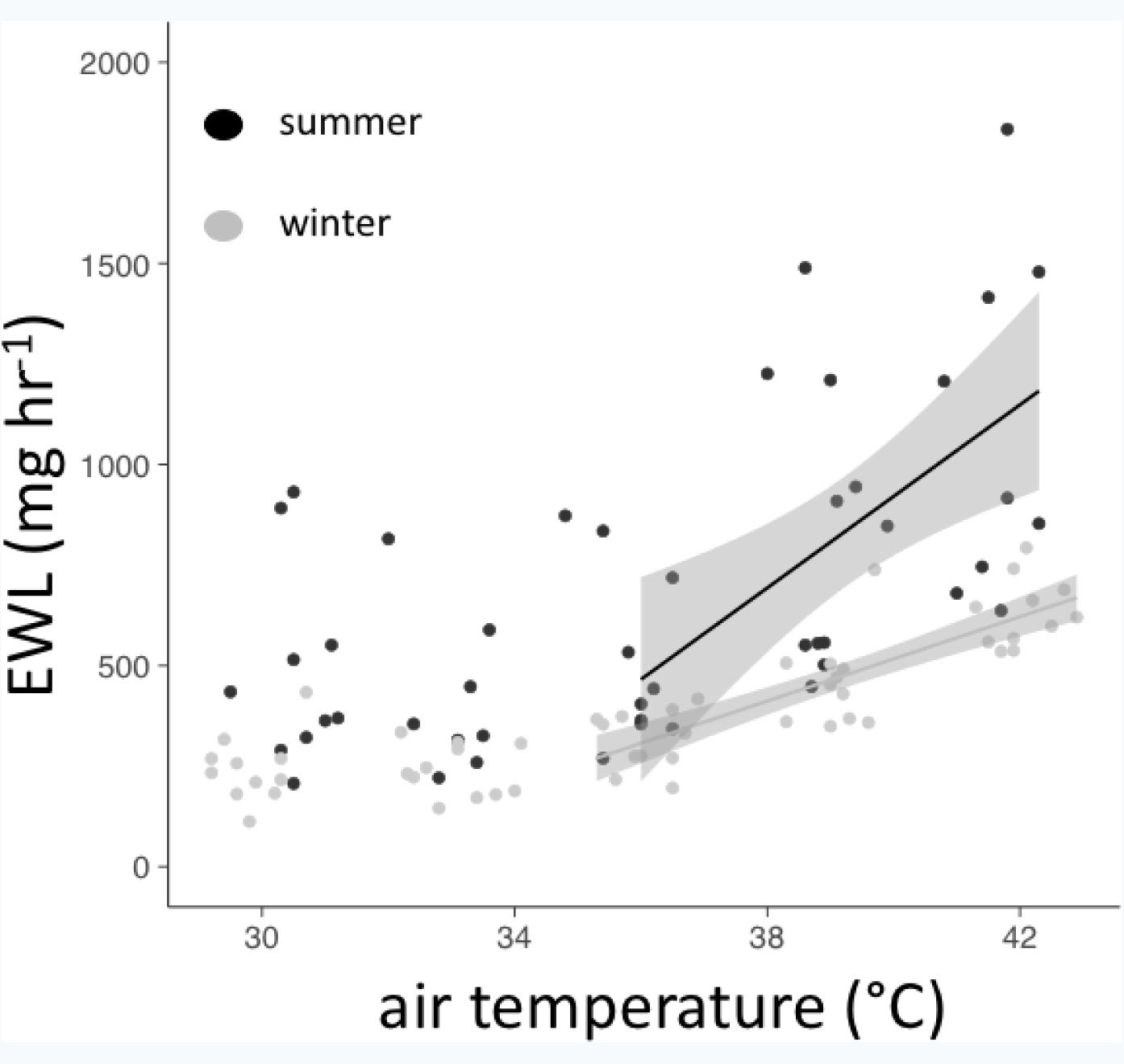
Evaporative water loss (mg hr^-1^) for Cape Rockjumpers (*Chaetops frenatus*) over a range of air temperatures (29 – 43 °C) captured in summer (n = 10) and winter (n = 11) at Blue Hill Nature Reserve, South Africa. Significant difference was found above an inflection point of 33.5 °C, with trendlines indicating best model fit above inflection points and 95% CI for each season.

### Resting Metabolic Rate

There were no significant inflection points for the relationship between RMR and T_air_ in either season. Despite increased EWL in summer we did not observe systematically higher summer RMR (W) at high T_air_. The only variable maintained in the best model explaining RMR was T_air_, where a significant increase in RMR was observed with increasing T_air_ (slope 0.02 ± 0.00; χ^2^_2,67_ = 26.38; p < 0.01; Fig 3; see supplementary materials Table S4). Any seasonal influence may have been masked by a higher variance in summer values compared to winter (mean RMR (W) ± SD: summer = 1.11 ± 0.46, winter = 0.87 ± 0.16).

**Figure 3.**
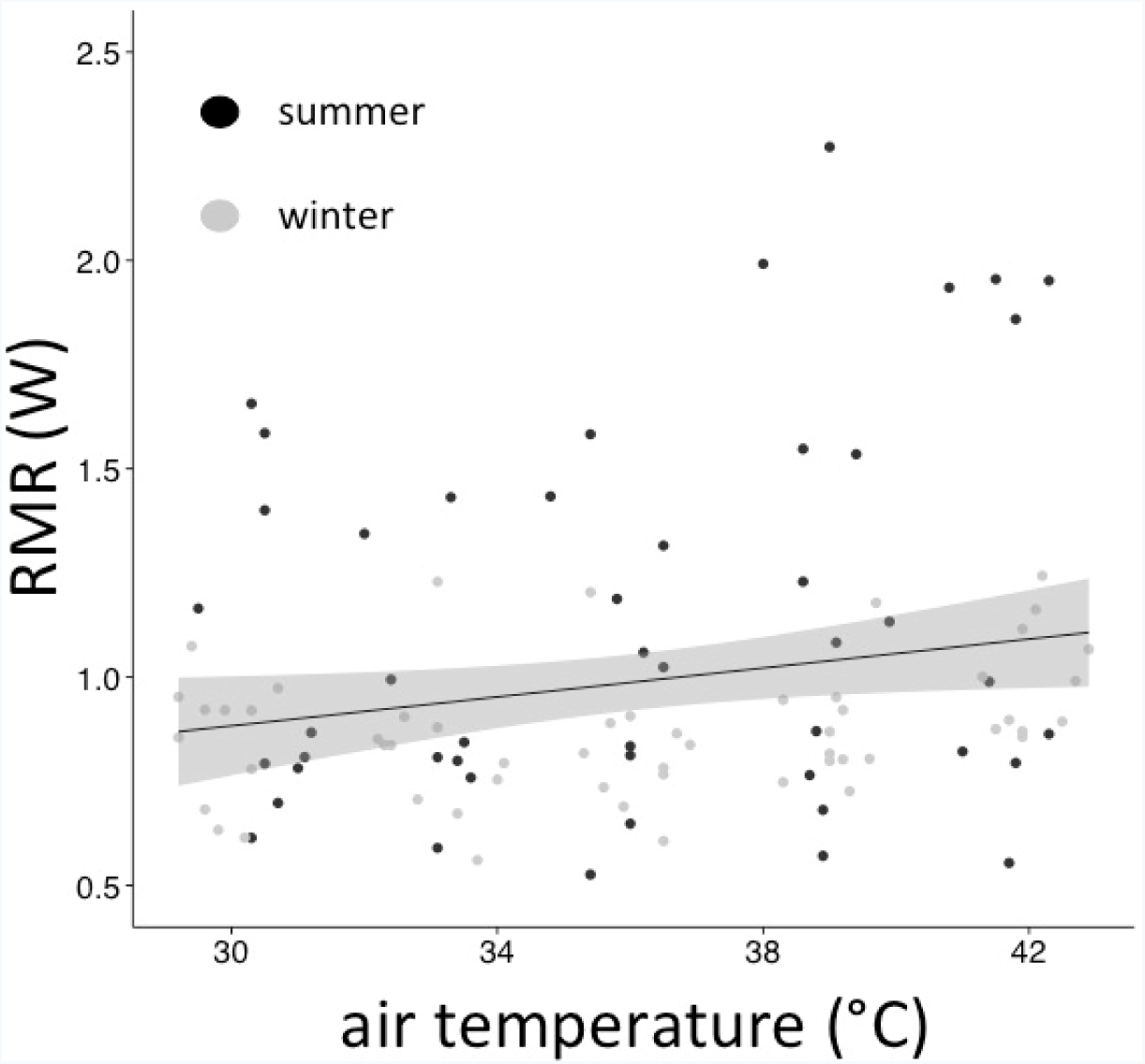
Resting metabolic rate (W) data over a range of air temperatures (29 – 43 °C) for Cape Rockjumpers (*Chaetops frenatus*) captured in summer (n = 10) and winter (n = 11) at Blue Hill Nature Reserve, South Africa. There were no significant differences between seasons with the trendline indicative of best model fit with 95% CI for both seasons combined.

### Body Temperature

The calculated T_b_ (°C) inflection point was 34.8 ± 2.0 °C in summer, with no significant inflection point found in winter. Despite the different inflection points between seasons, T_air_*season was not included in our top model, with the only significant effect on T_b_ that of increasing T_air_ (mean slope ± SD: summer = 0.30 ± 0.04 °C °C^-1^; winter = 0.14 ± 0.01 °C °C^-1^; χ^2^_1,63_ = 10.47; p < 0.01; Fig 4; see supplementary materials Table S5).

**Figure 4.**
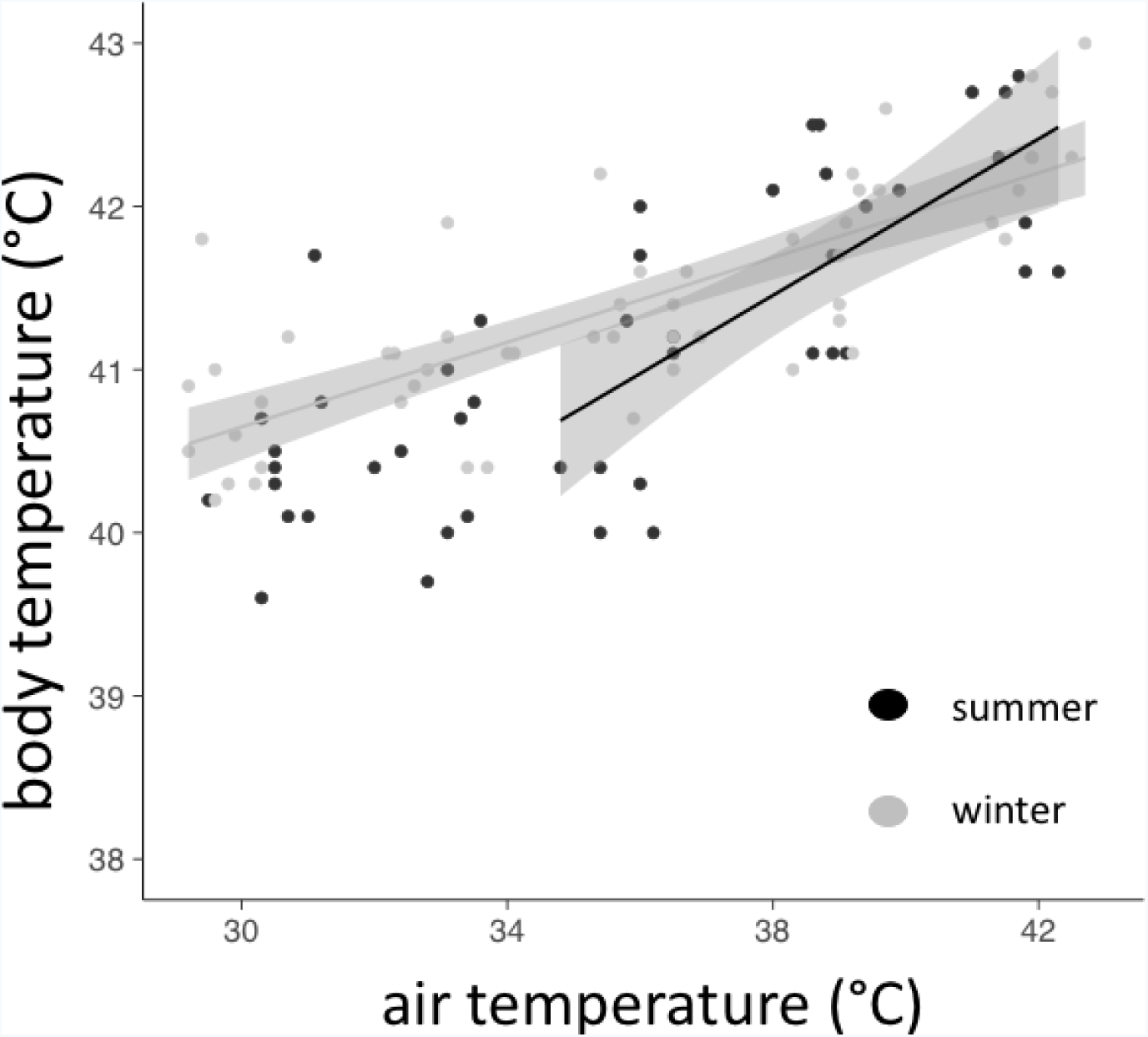
Body temperature (°C) data over a range of air temperatures (29 – 43 °C) for Cape Rockjumpers (*Chaetops frenatus*) captured in summer (n = 10) and winter (n = 11) at Blue Hill Nature Reserve, South Africa. No significant difference in season was found above an inflection point of 34.3°C in summer, with trendlines indicating best model fit above inflection point and 95% CI for each season.

**Table 1.** Evaporative cooling efficiency (evaporative heat loss [EHL] / metabolic heat production [MHP]), evaporative water loss (EWL), resting metabolic rate (RMR), and body temperature (T_b_), at T_air_ ≈ 30 and 42 °C from wild-living Cape Rockjumpers (*Chaetops frenatus*) captured during summer 2016 and winter 2015 at Blue Hill Nature Reserve, SouthAfrica. Data presented as mean ± SD. One female was excluded from presented data as she was removed from the chamber above T_air_ ≈ 39°C leaving a sample size of n = 9 in summer and n = 11 in winter.

|  | Summer<sup>i</sup> | Winter<sup>ii</sup> |
| --- | --- | --- |
| **EHL/MHP**<br>**T<sub>air</sub> ≈ 30 °C** | **.31 ± .08** | **.19 ± .04** |
| **EWL (mg hr<sup>-1</sup>)**<br>**T<sub>air</sub> ≈ 30 °C** | **526.46**<br>**± 266.50** | **243.63**<br>**± 83.55** |
| **RMR (W)**<br>**T<sub>air</sub> ≈ 30 °C** | **1.04 ± .38** | **.85 ± .15** |
| **T<sub>b</sub> (°C)**<br>**T<sub>air</sub> ≈ 30 °C** | **40.4 ± .6** | **40.7 ± .5** |
| **EHL/MHP**<br>**T<sub>air</sub> ≈ 42 °C** | **.58 ± .13** | **.42 ± .03** |
| **EWL (mg hr<sup>-1</sup>)**<br>**T<sub>air</sub> ≈ 42 °C** | **1085.13**<br>**± 418.60** | **631.52**<br>**± 84.40** |
| **RMR (W)**<br>**T<sub>air</sub> ≈ 42 °C** | **1.30 ± .60** | **.99 ± .13** |
| **T<sub>b</sub> (°C)**<br>**T<sub>air</sub> ≈ 42 °C** | **42.5 ± .8** | **42.6 ± .6** |

<sup>i</sup>n = 9; <sup>ii</sup>n = 11

## Discussion

Despite the dry summer conditions, seasonal thermoregulatory adjustments to heat in Rockjumper were centered on elevating EWL rates in summer compared to winter, with no clear seasonal change in RMR or T_b_. Indeed, the lack of seasonal difference in RMR in summer compared to winter was unexpected, as a lower RMR in summer should help maintain a lower T_b_ if birds were acclimatized to summer heat (Tieleman and Williams 2000, Williams and Tieleman 2005). The higher summer evaporative cooling efficiency in Rockjumpers was facilitated by higher EWL rates, contrary to our predictions that seasonal EWL adjustment would be centred on water conservation. The seasonal patterns of EWL in Rockjumpers are somewhat similar to those found in other species, despite these occupying habitats that are much warmer during summer; species such as Houbara Bustards (Tieleman et al. 2002), Freckled Nightjars (O’Connor et al. 2016) and mesic populations of Sparrow-weavers (Noakes et al. 2016) also elevate summer water demands.

We expected adjustments in metabolic heat production to play an important role in seasonal heat balance (Noakes et al. 2016), but this did not have an over-riding effect on evaporative efficiency in the Rockjumpers we studied. While we did not report a statistical difference in seasonal RMR, some individuals had RMR nearly two-fold higher than average (Table 1). We do not know the reasons for this variation among individuals. One potential reason is the fact that birds were in various stages of breeding; breeding activities were still underway in mid-summer when birds were captured for experiments (KN Oswald, personal observation). Yet, these individuals also had compensatory elevations in EWL and thus evaporative cooling efficiency was higher in summer.

The elevated EWL rates of Rockjumpers in summer compared to winter were in direct contrast to summer water conservation patterns observed in arid-zone Sparrow-weavers (Noakes et al. 2016). However, as with the present study, two mesic populations of Sparrow-weavers (Noakes et al. 2016) and a mesic population of Nightjars (O’Connor et al. 2016) increased EWL in summer compared to winter. We argue that the summer elevations in EWL should be more feasible in the above-mentioned mesic populations of Sparrow-Weavers and Nightjars compared to Cape Rockjumpers, given the differences in seasonal water bottlenecks we expect in their respective ranges.

The Sparrow-Weavers and Nightjars occupy a summer rainfall region (in contrast to the winter rainfall range of Cape Rockjumper) where the availability of water rich food or surface water increase during summer, and elevated EWL will be balanced easily by higher water acquisition rates (Smit and McKechnie 2015). Based on the rate of EWL at T_air_ ≈ 42 °C, we predict that Rockjumpers resting in full shade on a hot day (T_air_ ≈ 42°C) would evaporate 2.6 % M_b_ in water per hour. These values are quantitatively similar (average = 2.2 – 4.0 %) to those for Sparrow-weavers in both mesic and arid environments (Noakes et al. 2016). However, Sparrow-weavers that do not have access to surface water sources can balance their water intake by eating water rich food, particularly during wet summer months, when maximum T_air_s approach 40°C (Smit and McKechnie 2015, Smit et al. 2016). As of yet Rockjumpers have not been recorded drinking (Lee et al. 2017), and will need to balance water loss by obtaining water rich food on hot days. A better understanding of water intake rates at increasing temperatures are needed to properly predict dehydration risks in Rockjumpers under the dry summer conditions they experience naturally.

The average summer EHL/MHP ratio at T_air_ ≈ 42 °C was slightly higher in summer compared to winter (Table 1), but considerably lower than the average for Sparrow-weavers (summer = 1.32, winter = 1.04) at the same T_air_ (Noakes et al. 2016). This means at T_air_ ≈ T_b_ Sparrow-weavers dissipated more than 100% of their metabolic heat evaporatively, compared to only ~50% for Rockjumpers. This suggests Rockjumpers face a net heat gain at T_b_ ≈ T_air_, and the short time frame of our experiment at the maximum T_air_ may have precluded us from detecting a significant rise in T_b_.

Seasonal patterns in Rockjumper T_b_ regulation were also contrary to expected patterns of lower T_b_s at high T_air_ in summer, with T_b_ patterns suggesting Rockjumpers do not show seasonal differences in T_b_ patterns at hot temperatures. During both seasons T_b_ increased significantly as T_air_ increased from 34.2 °C to 42 °C, suggesting that Rockjumpers do not regulate a constant set-point T_b_ under hot conditions. Therefore it is unlikely we tested Rockjumpers near their thermal endpoint (Whitfield et al. 2015). Our experiment was not designed to determine thermal endpoints, but rather to assess seasonal changes in physiological costs at slightly higher T_air_s than Rockjumpers encounter naturally (maximum T_air_ at Blue Hill Nature Reserve 2012-2016 = 38.0 °C).

Rockjumper T_b_ at T_air_ ≈ 42 °C were quantitatively similar to other similar-sized passerines (e.g. Sparrow-Weaver average T_b_: summer = 42.0 °C, winter = 42.4 °C). However, the benefits, if any, of elevating T_b_ for Rockjumpers are not clear. Many authors have suggested that allowing T_b_ to rise above normothermic levels is a mechanism for storing metabolic heat (i.e. facultative hyperthermia) to combat risks of dehydration stemming from evaporative cooling (Tieleman and Williams 1999, Wolf 2000, Smit et al. 2013). The adaptive benefit of facultative hyperthermia is generally centred on water savings from decreasing EWL (e.g. (Maloney and Dawson 1998), which were not found for Rockjumpers. However, our emphasis on seasonal variation in thermoregulation as emphasis of acclimatization does not account for changes in breeding condition, changes in diet, or changes in water availability.

## Conclusion

Past studies have identified a number of traits that will Cape Rockjumpers vulnerable to climate change. These include a relatively small and declining climatic space, a fragmented range, overall low abundance (Lee and Barnard 2015a), recent population declines linked to warming climate, and a study showing low T_air_ inflections for increasing EWL in Rockjumpers compared to other similar sized species (Milne et al. 2015). Milne et al. (2015) argued low T_air_ inflections substantially elevate the costs of evaporative cooling in warmer parts of the species’ range, potentially providing a causal explanation for why Rockjumper population declines are greater in parts of their range where mean annual temperatures are rising.

Our study on seasonal physiological responses to heat show that Cape Rockjumpers have elevated water demands during the hot dry season of the year. If these physiological responses result in a seasonal water bottleneck our findings may partially explain declining populations in species with a restricted and well-defined climatic niche. However, to fully understand the effects on water budgets resulting from elevated EWL demands will require a more mechanistic approach such as was attempted for Sparrow-weavers (Smit and McKechnie 2015) and Night Parrots (*Pexoporus occidentalis*; (Kearney et al. 2016). Indeed, for Rockjumpers a modeling approach using closely related, more common species, may be the only alternative. Finally, our findings reiterate that avian seasonal physiological adjustments to heat may be as diverse as adjustments to cold. Seasonal studies on thermoregulation in the heat will greatly improve our knowledge of the functional (or adaptive) value traits such evaporative cooling efficiency and heat tolerance hold and how they contribute to the physiological stress organisms experience in heterogenous environments.

## Data Accessibility

Our raw data has been made accessible as an online file “Table S10.csv” under the associated supplementary materials.

## Acknowledgments

We firstly thank the Lee family for allowing us to conduct research on their property. We would also like to thank the volunteers who spent many hours helping us catch Rockjumpers: Audrey Miller, Jenny Tartini, Alacia Welch, Gavin Emmons, Cristina Ebneter, Nicolas Pattinson, Cuen Muller, and Maxine Smit. Special thanks to Mark Brigham, and two anonymous reviewers for commenting on drafts of this manuscript. This study was funded by a National Research Foundation Thuthuka Grant (BS) and a Nelson Mandela Metropolitan University Research Themes Grant (BS). All experimental procedures were approved by the Research Ethics Committee: Animal (A15-SCI-ZOO-007) at Nelson Mandela Metropolitan University with bird capture permit issued by Cape Nature, Western Cape, South Africa (0037-AAA041-00060).

## References

Baker, W. C., and J. F. Pouchot. 1983. The measurement of gas flow part ii. Journal of the Air Pollution Control Association33:156–162.

Barton, K. 2013. MuMIn: Multi-model inference. R package version 1.9. 5. R Project for Statistical Computing, Vienna, Austria.

Bates, D., M. Maechler, B. Bolker, and S. Walker. 2013. lme4: Linear mixed-effects models using Eigen and S4. R package version 1

CooperS. J. 2002. Seasonal metabolic acclimatization in mountain chickadees and juniper titmice. Physiol. Biochem. Zool. 75:386–395.

Cowling, R. M., A. J. Potts, P. L. Bradshaw, J. Colville, M. Arianoutsou, S. Ferrier, F. Forest, N. M. Fyllas, S. D. Hopper, and F. Ojeda. 2015. Variation in plant diversity in mediterranean climate ecosystems: the role of climatic and topographical stability. J. Biogeogr. 42:552–564.

Fox, J., G. G. Friendly, S. Graves, R. Heiberger, G. Monette, H. Nilsson, B. Ripley, S. Weisberg, M. J. Fox, and M. Suggests. 2007. The car package.

Gerson, A. R., E. K. Smith, B. Smit, A. E. Mckechnie, and B. O. Wolf. 2014. The impact of humidity on evaporative cooling in small desert birds exposed to high air temperatures. Physiol. Biochem. Zool. 87:782–795.

Gibbons, W. J., and K. M. Andrews. 2004. PIT tagging: simple technology at its best. Bioscience54:447–454.

Kearney, M. R., W. P. Porter, and S. A. Murphy. 2016. An estimate of the water budget for the endangered night parrot of Australia under recent and future climates. Climate Change Responses3:14.

Lee, A., and P. Barnard. 2015a. Endemic birds of the Fynbos biome: a conservation assessment and impacts of climate change. Bird Conservation International 1:1–17.

Lee, A., and P. Barnard. 2015b. Endemic birds of the Fynbos biome: a conservation assessment and impacts of climate change. Bird Conservation International:1–17.

Lee,A. T., D. Wright, and P. Barnard. 2017. Hot bird drinking patterns: drivers of water visitation in a fynbos bird community. Afr. J. Ecol.

LightonJ. R. 2008. Measuring metabolic rates: a manual for scientists. Oxford University Press.

Maloney, S. K., and T. J. Dawson. 1998. Changes in pattern of heat loss at high ambient temperature caused by water deprivation in a large flightless bird, the emu. Physiol. Zool. 71:712–719.

Marder, J., and Y. Arieli. 1988. Heat balance of acclimated pigeons (Columba livia) exposed to temperatures up to 60 C Ta. Comparative Biochemistry and Physiology Part A: Physiology91:165–170.

Marras, S., A. Cucco, F. Antognarelli, E. Azzurro, M. Milazzo, M. Bariche, M. Butenschön, S. Kay, M. Di Bitetto, and G. Quattrocchi. 2015. Predicting future thermal habitat suitability of competing native and invasive fish species: from metabolic scope to oceanographic modelling. Conservation Physiology 3:cou059

Mckechnie, A. E., M. J. Noakes, and B. Smit. 2015. Global patterns of seasonal acclimatization in avian resting metabolic rates. Journal of Ornithology 156:367–376.

Mckechnie, A. E., and B. O. Wolf. 2004. Partitioning of evaporative water loss in white-winged doves: plasticity in response to short-term thermal acclimation. J. Exp. Biol. 207:203–210.

Milne, R., S. J. Cunningham, A. T. Lee, and B. Smit. 2015. The role of thermal physiology in recent declines of birds in a biodiversity hotspot. Conservation Physiology 3:cov048

Mucina, L., and M. C. Rutherford. 2006. The vegetation of South Africa, Lesotho and Swaziland. South African National Biodiversity Institute.

Muggeo V. M. 2008. Segmented: an R package to fit regression models with broken-line relationships. Pages 20–25 R news

Noakes, M. J., B. O. Wolf, and A. E. Mckechnie. 2016. Seasonal and geographical variation in heat tolerance and evaporative cooling capacity in a passerine bird. J. Exp. Biol. 219:859–869.

O’connor, T. 1995. Metabolic characteristics and body composition in house finches: effects of seasonal acclimatization. Journal of Comparative Physiology B 165:298–305.

O’connor, R. S., B. O. Wolf, R. M. Brigham, and A. E. Mckechnie. 2016. Avian thermoregulation in the heat: efficient evaporative cooling in two southern African nightjars. Journal of Comparative Physiology B:1–15.

Ophir, E., Y. Arieli, J. Marder, and M. Horowitz. 2002. Cutaneous blood flow in the pigeon Columba livia: its possible relevance to cutaneous water evaporation. J. Exp. Biol. 205:2627–2636.

Oswald, K. N., A. A. Evlambiou, Â. M. Ribeiro, and B. Smit. 2017. Tag location and risk assessment for PIT tagging passerines. Ibis.

Prinzinger, R., A. Pressmar, and E. Schleucher. 1991. Body temperature in birds. Comparative Biochemistry and Physiology Part A: Physiology 99:499–506.

Ratnayake, C. P., C. Morosinotto, S. Ruuskanen, A. Villers, and R. L. Thomson. 2014. Passive integrated transponders (PIT) on a small migratory passerine bird: absence of deleterious short and long-term effects. Ornis Fenn. 91

Smit, B., C. Harding, P. Hockey, and A. E. Mckechnie. 2013. Adaptive thermoregulation during summer in two populations of an arid-zone passerine. Ecology 94:1142–1154.

Smit, B., and A. E. Mckechnie. 2015. Water and energy fluxes during summer in an arid zone passerine bird. Ibis 157:774–786.

Smit, B., G. Zietsman, R. Martin, S. Cunningham, A. Mckechnie, and P. Hockey. 2016. Behavioural responses to heat in desert birds: implications for predicting vulnerability to climate warming. Climate Change Responses 3:9.

Swanson, D. L. 1991. Seasonal adjustments in metabolism and insulation in the dark-eyed junco. Condor:538–545.

Swanson, D. L. 2010. Seasonal metabolic variation in birds: functional and mechanistic correlates. Current Ornithology 17:75–129.

Tieleman B. I. 2002. Avian adaptation along an aridity gradient: physiology, behavior, and life history. [sn]

Tieleman, B. I., J. B. Williams. 1999. The role of hyperthermia in the water economy of desert birds. Physiol. Biochem. Zool. 72:87–100.

Tieleman, B. I., J. B. Williams. 2000. The adjustment of avian metabolic rates and water fluxes to desert environments. Physiol. Biochem. Zool. 73:461–479.

Tieleman, B. I., J. B. Williams. 2002. Cutaneous and respiratory water loss in larks from arid and mesic environments. Physiol. Biochem. Zool. 75:590–599.

Tieleman, B. I., J. B. Williams, F. Lacroix, and P. Paillat. 2002. Physiological responses of Houbara bustards to high ambient temperatures. J. Exp. Biol. 205:503–511.

Walsberg, G., and B. Wolf. 1995. Variation in the respiratory quotient of birds and implications for indirect calorimetry using measurements of carbon dioxide production. The Journal of experimental biology 198:213–219.

Whitfield, M. C., B. Smit, A. E. Mckechnie, and B. O. Wolf. 2015. Avian thermoregulation in the heat: scaling of heat tolerance and evaporative cooling capacity in three southern African arid-zone passerines. J. Exp. Biol. 218:1705–1714.

Williams, J. B., and B. I. Tieleman. 2005. Physiological adaptation in desert birds. Bioscience 55:416–425.

Wolf B. 2000. Global warming and avian occupancy of hot deserts; a physiological and behavioral perspective. Rev. Chil. Hist. Nat. 73:395–400.

Zuur, A., E. N. Ieno, N. Walker, A. A. Saveliev, and G. M. Smith. 2009. Mixed effects models and extensions in ecology with R. Springer.

